# New Segmentation and Feature Extraction Algorithm for Classification of White Blood Cells in Peripheral Smear Images

**DOI:** 10.1101/2021.04.29.441751

**Authors:** Sajad Tavakoli, Ali Ghaffari, Zahra Mousavi Kouzehkanan, Reshad Hosseini

## Abstract

This article addresses a new method for the classification of white blood cells (WBCs) using image processing techniques and machine learning methods. The proposed method consists of three steps: detecting the nucleus and cytoplasm, extracting features, and classification. At first, a new algorithm is designed to segment the nucleus. For the cytoplasm to be detected, only a part of it located inside the convex hull of the nucleus is involved in the process. This attitude helps us overcome the difficulties of segmenting the cytoplasm. In the second phase, three shapes and four novel color features are devised and extracted. Finally, by using an SVM model, the WBCs are classified. The segmentation algorithm can detect the nucleus with a dice similarity coefficient of 0.9675. The proposed method can categorize WBCs in Raabin-WBC, LISC, and BCCD datasets with accuracies of 94.65 %, 92.21 %, and 94.20 %, respectively. It is worth mentioning that the hyperparameters of the classifier are fixed only with the Raabin-WBC dataset, and these parameters are not readjusted for LISC and BCCD datasets. The obtained results demonstrate that the proposed method is robust, fast, and accurate.

## Introduction

Generally, there exist three types of blood cells: red blood cells, white blood cells (WBC), and platelets; among these, WBCs are responsible for the immune system and protect the body against diseases and infections. In peripheral blood, WBCs are categorized into five general types: lymphocytes, monocytes, neutrophils, eosinophils, and basophils. In various diseases such as leukemia, anemia, malaria, human immunodeficiency virus infection (HIV) and, infectious diseases, changes in the number of WBCs are visible [1, 2, 3, 4]. A recent study also indicated that leukopenia, lymphocytopenia, and eosinophil cytopenia have occurred significantly more in Covid-19 patients [5]. Therefore, differential counting of WBCs can be of considerable assistance in disease diagnosis [6].

In several cases, peripheral blood smear analysis is done manually by a hematologist who visually analyzes the blood smears under the microscope [1, 7]. This procedure is very time consuming [8] and can be inaccurate due to tiredness and human error [9]. On the other hand, automated hematology analyzer devices (e.g. Sysmex) are very expensive, especially for low-income countries [10]. Fortunately, machine learning-based methods can easily fill the above-mentioned gaps. This study is aimed at proposing a new method based on machine learning and image processing techniques to classify WBCs in peripheral blood smear.

In machine learning-based methods, at first, it is a requisite to collect the appropriate dataset taking quality, variety, and size into account. Yet, the lack of such dataset with aforementioned properties is the major challenge [11]. A plethora of articles have used small datasets collected using only one microscope or one camera [7, 12, 13]. Also, some of these datasets were solely labeled by one hematologist [7, 14], which carries the risk of being labeled incorrectly because of the challenges diagnosing WBC types involves. In this study, three different datasets were used: Raabin-WBC dataset [15], LISC dataset [7], and BCCD dataset [16]. We will elaborate more on these datasets in the datasets section.

After data collection, diverse machine learning techniques can be used to classify WBCs. In general, different methods proposed in the literature for classifying WBCs favor either traditional or deep learning frameworks [1]. In traditional frameworks, it is first necessary to extract the appropriate handcraft features from WBCs, and then, classify them using one or an ensemble of several classifiers. Feature engineering is the most challenging part of traditional approaches. Unlike traditional frameworks, in deep learning frameworks, features are automatically extracted by means of deep neural networks.

One of the most commonly used networks for classifying images are convolutional neural networks (CNNs). To obtain good classification results, we need a large deep CNN with numerous parameters. Training such a large network from scratch needs a large dataset. However, medical datasets are not usually large enough. Therefore pre-trained networks are normally used in two ways. The first way is to extract features by means of a pre-trained network as the input of a traditional classifier model such as support vector machine (SVM), k nearest neighbor (KNN), etc. The second way is to fine-tune the pre-trained network using a small dataset..

There are some works that have utilized pre-trained CNNs for extracting features in the task of classifying WBCs [17, 18, 19, 20]. In the task of diagnosing acute lymphoblastic leukemia, Rehman *et al*. [17] compared the accuracy of using three different classifiers on the image features extracted by pre-trained CNNs. They observed that the SVM classifier gives the best results. In [18], the features extracted by three well known CNN architectures (AlexNet, GoogleNet, and ResNet-50) were merged, then proper features were selected using the maximal information coefficient and ridge algorithms. Finally, WBCs were classified using a quadratic discriminant analysis model. Similarly, Togacar *et al*. [19] used a pre-trained Alexnet network to extract features and a quadratic discriminant analysis model to classify WBCs. Sahlol *et al*. [20] employed a pre-trained CNN for extracting features, along with a statistically enhanced salp swarm algorithm for feature selection, and an SVM model.

Deep learning neural networks can also be directly trained to categorize WBCs [1, 21, 22, 23, 24, 25, 26]. Hedge *et al*. [1] performed the classification of WBCs with and without using a pre-trained network. They found out that full training from scratch leads to better results than fine-tuning an AlexNet pre-trained network. In [21], authors addressed the classification of WBCs by tuning pre-trained AlexNet and LeNet-5 networks as well as training a new CNN from scratch. They declared that the novel network they have proposed performed better than the fine-tuned networks mentioned previously. Jung et al. [22] designed a new CNN architecture called W-Net to classify WBCs in the LISC dataset. Baydilli and Atila [23] adopted capsule networks to classify the WBCs existing in the LISC dataset. Banik et al. [24] devised a fused CNN model in the task of differential WBC count and evaluated their model with the BCCD dataset. Liang *et al*. [25] combined the output feature vector of the flatten layer in a fine-tuned CNN and a long short term memory network to classify WBCs in BCCD dataset. A new complicated fused CNN introduced in [26] was trained from scratch on 10253 augmented WBCs images from the BCCD dataset. Despite the complexity of the proposed CNN in [26], the number of its parameters stands at 133000.

For the classification of WBCs based on traditional frameworks, segmenting the nucleus and the cytoplasm of WBCs is a vital but tough task. In this study, a novel accurate method to segment the nucleus is put forward. In order to segment the nucleus, some researchers used the thresholding algorithms after applying various pre-processing techniques on the image (e.g. Otsu’s thresholding algorithm, Zack algorithm, and etc.) [27, 28, 29, 30]. A combination of machine learning and image processing techniques is also commonly employed to segment the nucleus of the WBC [31, 32]. Moreover, during the last decade, CNNs have gained more popularity and are used to segment the nucleus of the WBC and cytoplasm [33]. Segmenting the cytoplasm is more complicated and less accurate than segmenting the nucleus. Therefore, in this paper, a part of the cytoplasm rather than the whole cytoplasm is detected as a representative of the cytoplasm (ROC) to be segmented. This approach, as a result, does not have the difficulties of segmenting the cytoplasm. We will talk more about this method in materials and methods section.

In order to classify WBCs after segmenting the nucleus and the cytoplasm, discriminative features need to be extracted. Shape characteristics such as circularity, convexity, solidity are meaningful features for the nucleus. This is due to the fact that lymphocytes and monocytes are mononuclear, and the shape of their nucleus is circular and ellipsoidal, respectively [11]. On the other hand, the nucleus of neutrophil and eosinophil is multi-lobed [11] and non-solid. Characteristics such as color and texture, e.g., local binary pattern (LBP) or gray level co-occurrence matrix (GLCM), are also interpretable features for the cytoplasm [11]. In addition to the mentioned features, SIFT (scale-invariant features transform) or dense SIFT algorithm can be employed for feature extraction. In the next paragraph, we review some related works that use traditional frameworks for classifying WBCs.

Rezatofighi and Soltani-zadeh [7] proposed a new system for the classification of five types of WBCs. In this system, nucleus and cytoplasm were extracted using the Gram-Schmidt method and Snake algorithm, respectively. Then, LBP and GLCM were used for feature extraction, and WBCs were categorized using a hybrid classifier including a neural network and an SVM model. Hiremath *et al*. [28] segmented the nucleus utilizing a global thresholding algorithm and classified WBCs using geometric features of the nucleus and cytoplasm. In [29], Otsu’s thresholding algorithm was used to detect the nucleus, and shape features such as area, perimeter, eccentricity, and circularity were extracted to identify five types of WBCs. Diagnosing ALL using images of WBCs was investigated in [30]. The authors of this paper applied the Zack algorithm to estimate the threshold value to segment the cells. Then, shape, texture, and color features were extracted, and the best features were selected by the means of the social spider optimization algorithm. Finally, they classified WBCs into two types of healthy and non-healthy, using a KNN classifier. Ghane *et al*. [31] designed a new method to segment the nucleus of the WBCs through a novel combination of Otsu’s thresholding algorithm, k-means clustering, and modified watershed algorithm, and succeeded in segmenting nuclei with a precision of 96.07 %. Laosai and Chamnongthai [32] examined the task of diagnosing ALL and acute myelogenous leukemia using the images of the WBCs. They detected the nuclei by employing the k-means clustering algorithm, extracted shape and texture features, and finally categorized WBCs utilizing an SVM classifier.

In this section, we briefly introduced the WBCs, its clinical importance and available datasets together with methods used to classify and count WBCs in other studies. In the materials and methods section, we present our proposed method for classifying WBCs. Afterwards, we will present and compare the obtained results with those of the other studies.

## Materials and Methods

### Datasets

Three different datasets used in this study are Raabin-WBC [15], LISC [7], and BCCD [16]. These datasets are discussed in the next three subsections, and are compared in Table 1. Also, Figure 1 shows some sample images of these three datasets.

**Table 1.** The properties of LISC, BCCD, and Raabin-WBC datasets.

| Dataset |  | Number of WBCs |  |  |  |  |  | Staining | Microscope and Zoom | Camera |
| --- | --- | --- | --- | --- | --- | --- | --- | --- | --- | --- |
|  |  | Lymph. | Mon. | Neut. | Eos. | Bas. | Total |  |  |  |
| Raabin-WBC [15] | All WBCs | 3461 | 795 | 8891 | 1066 | 301 | 14514 | Giemsa | 1.Olympus CX18<br>2.Zeiss microscope<br>Zoom : 100X | 1.Phone camera-Samsung Galaxy S5<br>2.Phone camera-LG G3 |
|  | Training set | 2427 | 561 | 6231 | 744 | 212 | 10175 |  |  |  |
|  | Augmented training set | 7305 | 6083 | 6231 | 6680 | 3180 | 29479 |  |  |  |
|  | Test set | 1034 | 234 | 2660 | 322 | 89 | 4339 |  |  |  |
| BCCD [16] | All WBCs | 33 | 19 | 208 | 86 | 3 | 349 | Gismo-right | Regular light microscope<br>Zoom : 100X | CCD color camera |
|  | Training set | 27 | 16 | 159 | 75 | - | 277 |  |  |  |
|  | Augmented training set | 2483 | 2478 | 2499 | 2497 | - | 9957 |  |  |  |
|  | Test set | 6 | 3 | 49 | 11 | - | 72 |  |  |  |
| LISC [7] | All WBCs | 59 | 48 | 56 | 39 | 55 | 257 | Gismo-right | Axioskope40<br>Zoom : 100X | Sony-SSCDC50AP |
|  | Training set | 41 | 33 | 39 | 28 | 39 | 180 |  |  |  |
|  | Augmented training set | 410 | 396 | 390 | 420 | 390 | 2006 |  |  |  |
|  | Test set | 18 | 15 | 17 | 11 | 16 | 77 |  |  |  |

**Figure 1.**
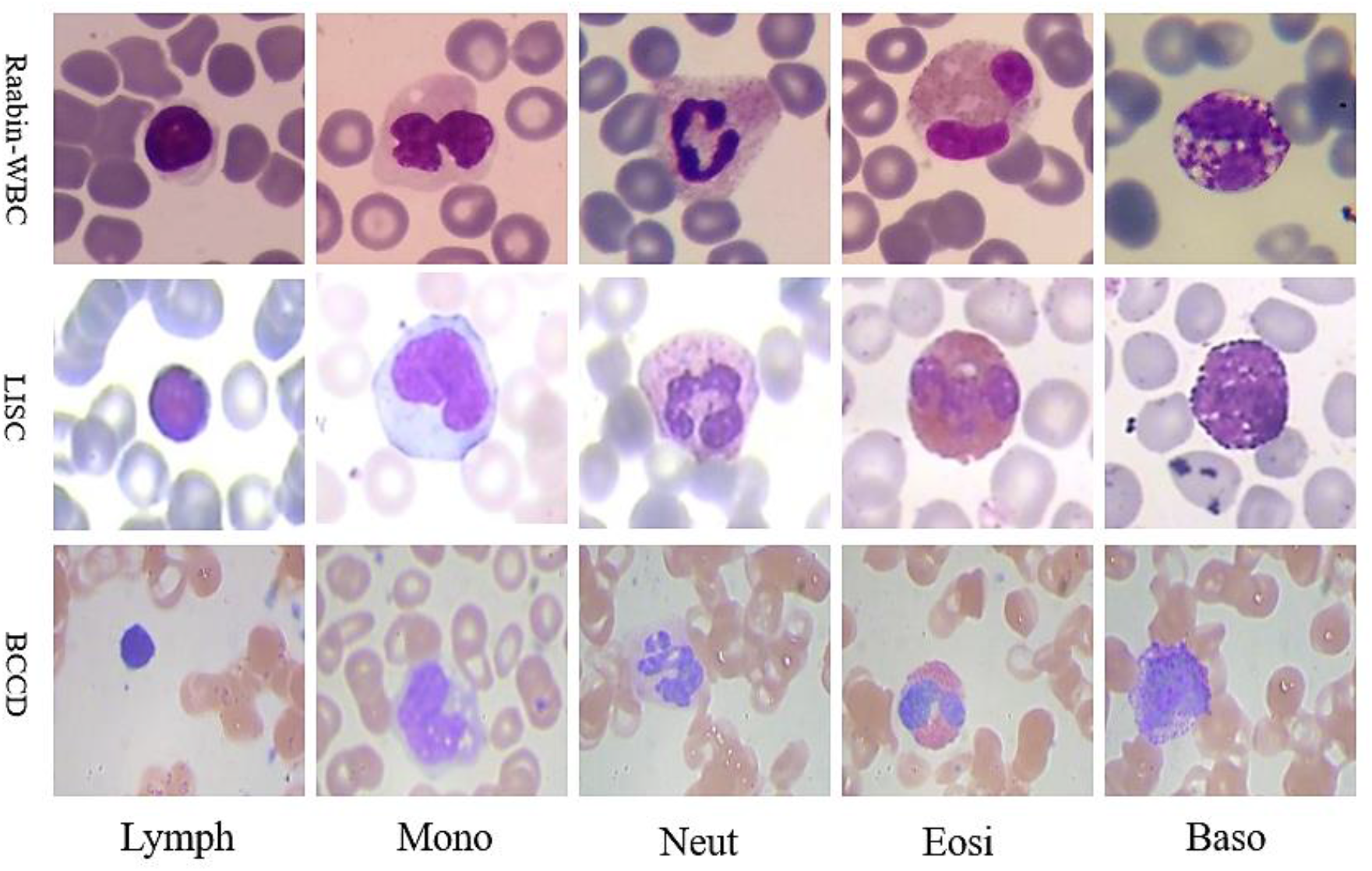
Some samples of the WBCs in Raabin-WBC, LISC, and BCCD datasets; Lymph (lymphocyte), Mono (monocyte), Neut (neutrophil), Eosi (eosinophil), Baso (basophil).

#### Raabin-WBC dataset

Raabin-WBC [15] is a large free-access dataset recently published in 2021. Raabin-WBC dataset possesses three sets of WBC cropped images for classification: Train, Test-A, and Test-B. All WBCs in Train and Test-A sets have been separately labeled by two experts. Yet, images of Test-B have not yet been labeled thoroughly. Therefore, in this study we only used Train and Test-A sets. These two sets have been collected from 56 normal peripheral blood smears (for lymphocyte, monocyte, neutrophil, and eosinophil) and one chronic myeloid leukemia (CML) case (for basophil) and contain 14514 WBC images. All these films were stained through Giemsa technique. The normal peripheral blood smears have been taken using the camera phone of Samsung Galaxy S5 and the microscope of Olympus CX18. Also, the CML slide has been imaged utilizing an LG G3 camera phone along with a microscope of Zeiss brand. It is worth noting that the images have all been taken with a magnification of 100.

#### LISC dataset

LISC dataset [7] contains 257 WBCs from peripheral blood, which have been labeled by only one expert. The LISC dataset has been acquired from peripheral blood smear and stained through Gismo-right technique. These images have been taken at a magnification of 100 using a light microscope (Microscope-Axioskope 40) and a digital camera (Sony Model No. SSCDC50AP). We cropped all WBCs in this dataset as shown in Figure 1.

#### BCCD dataset

BCCD dataset [16] has been taken from the peripheral blood and includes 349 WBCs labeled by one expert. The Gismo-right technique has been employed for staining the blood smears. This dataset, also, has been imaged at a magnification of 100 using a regular light microscope together with a CCD color camera [34]. In addition, based on diagnosis made by two of our experts, we found that one of the images of the BCCD dataset had been incorrectly labeled, and thus, we corrected this label.

#### Training, augmented training, and test sets

For the Raabin-WBC dataset, we have employed already split sets of the original data namely Train and Test-A sets for training and test. In this dataset, different blood smears have been considered for the training and testing sets. Test-A and Train sets comprise almost 30 percent and 70 percent of the whole data, respectively. For the LISC dataset, we randomly selected 70 percent of the data for training, and 30 percent for testing. BCCD dataset has two splits in the original data, 80% of which serve as training and 20% as testing. Since this dataset had only three basophils, we ignored the basophils in BCCD and only considered the remaining four types.

To train an appropriate classifier, it is necessary to balance the training data adopting various augmentation methods. For this reason, some augmentation methods such as horizontal flip, vertical flip, random rotation (between −90 and +90 degree), random scale augmentation (rescaling between 0.8 and 1.2), and a combination of them were utilized to augment the training sets of Raabin-WBC and LISC datasets. In addition, the training data of the BCCD dataset had already been augmented. In Table 1, all information about the amount of data in each set is presented.

### Nucleus Segmentation

Three following steps for nucleus segmentation are considered: Firstly, a color balancing algorithm [1] is applied to the RGB input image, then the CMYK and HLS color spaces are computed and combined and a soft map is computed. Finally, the nucleus is segmented by applying Otsu’s thresholding algorithm on the aforementioned soft map. The precise steps of the nucleus segmentation algorithm are as follows:

a. *Converting color-balanced RGB image to CMYK color space*
b. *KM = (K component) – (M component)*
c. *Converting color-balanced RGB image to HLS color space*
d. *MS = Min(M component, S component)*
e. *Output soft map = MS – KM*
f. *Employing Otsu’s thresholding algorithm to segment the nucleus*

Figures 2 illustrates the resulting images obtained by applying different steps of the proposed method to segment a sample nucleus. As depicted in Figure 2, red blood cells and the cytoplasm of the WBC in the K component have more intensity than those in the M component. Furthermore, the nucleus of the WBC has a lower intensity in comparison to the M component. Accordingly, as shown in Figure 2 (6), subtracting the M component from the K component produces an image the nucleus pixels of which are zero or close to zero. On the other hand, as seen in Figure 2 (7), computing the minimum of the M and S channels creates an image wherein the intensity of the red blood cell and the background are close to zero. Finally, by subtracting Figure 2 (6) from Figure 2 (7), Figure 2 (8) is formed in which the red blood cells, cytoplasm, and the background are eliminated. Figure 3 also shows the block-diagram of the proposed algorithm.

**Figure 2.**
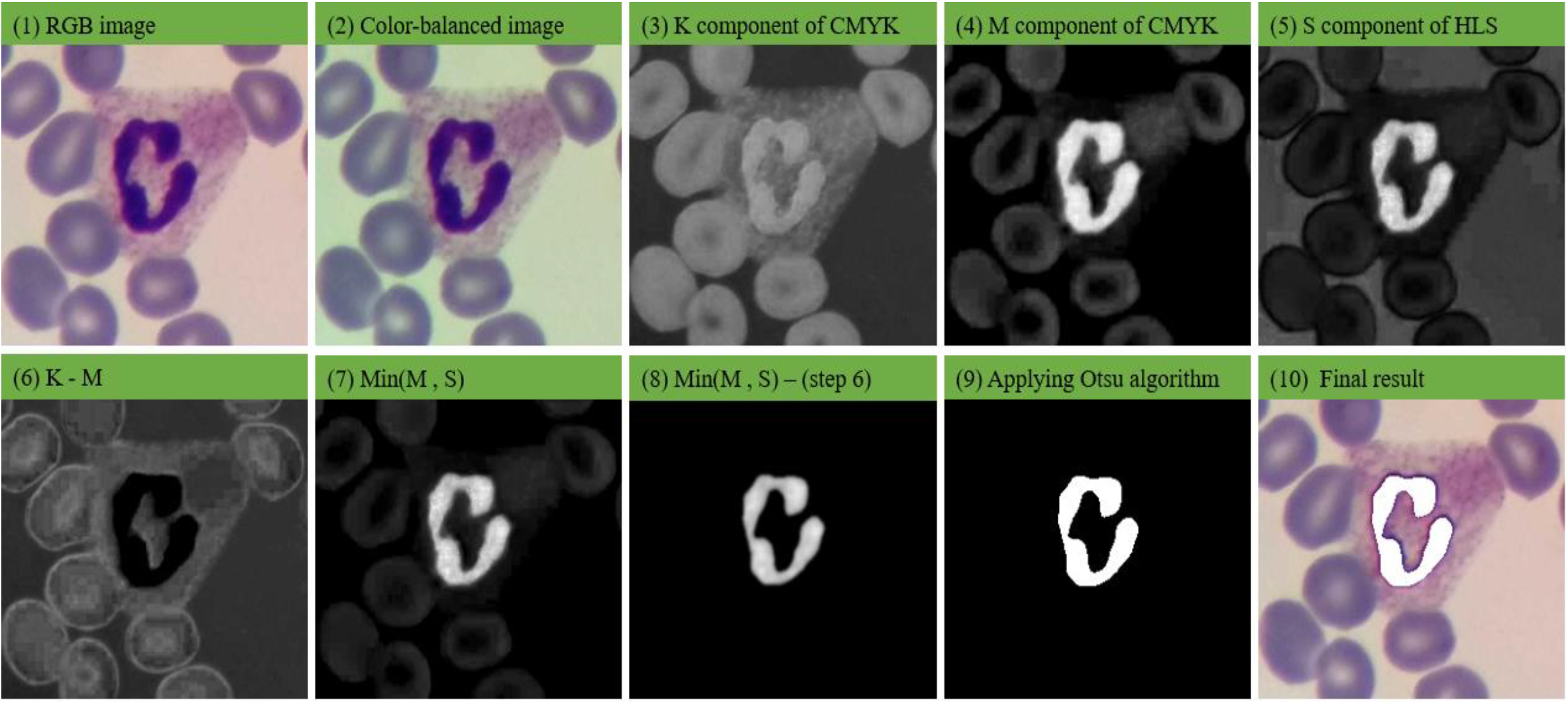
The results obtained through applying different steps of our nucleus segmentation method: (1) RGB image, (2) color-balanced image, (3) K component of CMYK color space, (4) M component of CMYK color space, (5) S component of HLS color space, (6) result of K – M, (7) result of Min(M, S), (8) result of Min(M, S) – (K – M), (9) the result of applying Otsu’s thresholding algorithm, (10) the final result.

**Figure 3.**
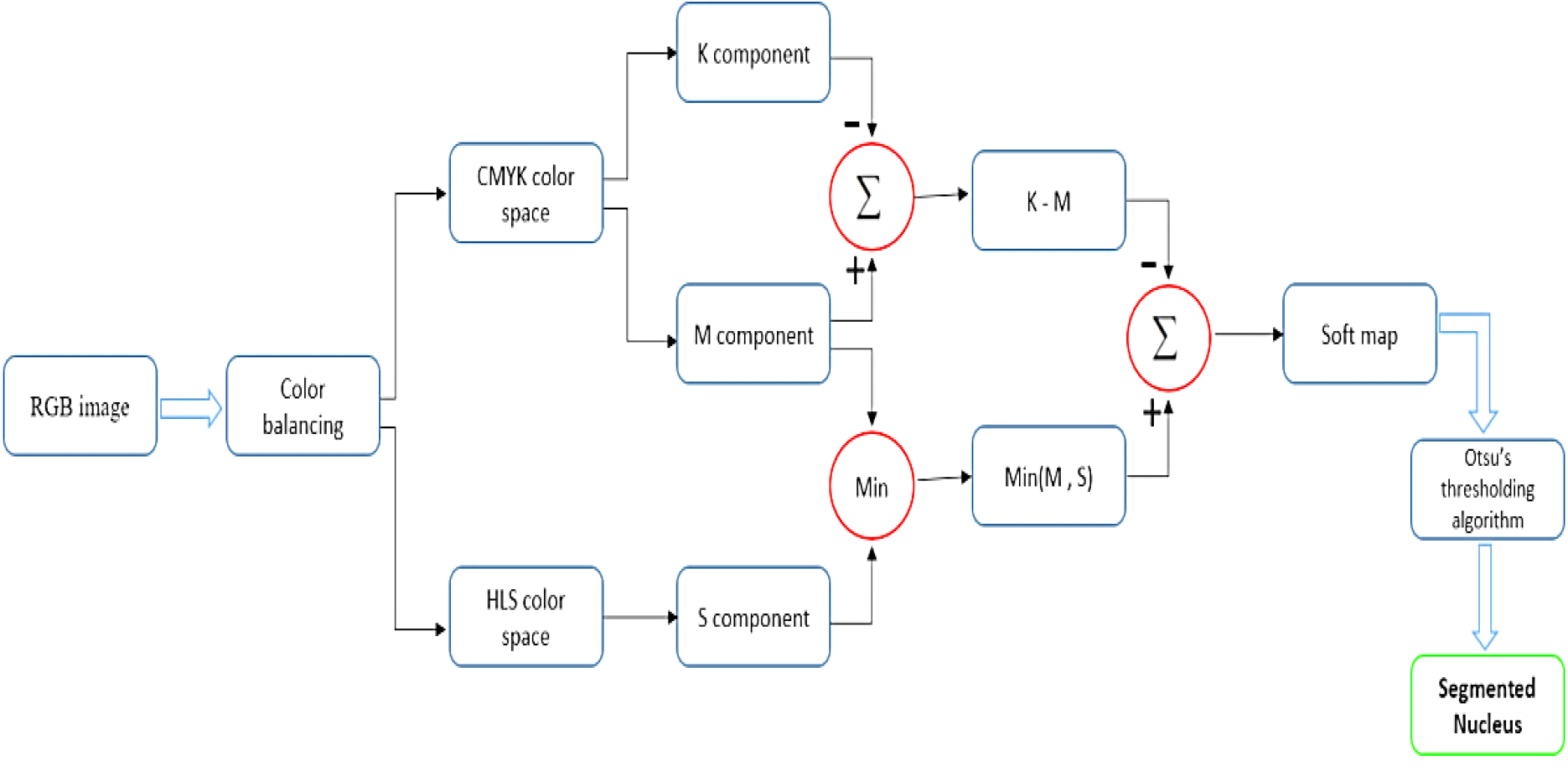
The block diagram of the nucleus segmentation method.

In this research, the color balancing algorithm of [1] is utilized to reduce color variations. To create a color-balanced representation of the image, it is necessary to compute the mean of R, G, and B channels as well as the grayscale representation of the RGB image. Then, by using equation (1), the new balanced R, G, B components are obtained.

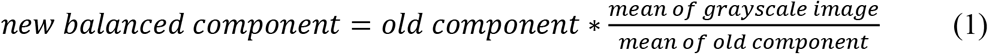

After color balancing, the color-balanced image is converted to the CMYK and HLS color spaces. The CMYK color space has four components, which are cyan (C), magenta (M), yellow (Y), and black (K). The steps of converting RGB to CMYK are given by the following equations [37]:

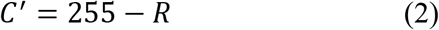

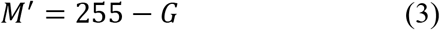

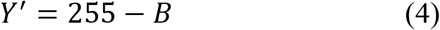

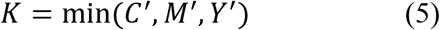

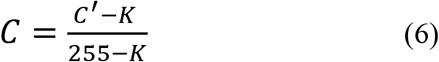

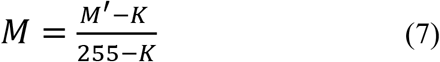

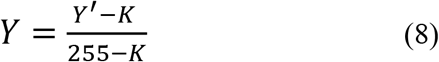

### Cytoplasm Detection

To extract proper features from the cytoplasm, it is first necessary to segment it. However, segmenting the cytoplasm is more difficult and less accurate than segmenting the nucleus. Hence, we designed a new method to solve this problem. In this method, the convex hull of the nucleus is obtained first, and a part of the cytoplasm that has been located inside the convex hull is considered as the representative of the cytoplasm (ROC). The more convex nucleus is, the smaller ROC is. Thus, lymphocytes, which usually have a circular nucleus, have lower ROC than neutrophils. Figure 4 illustrates this point.

**Figure 4.**
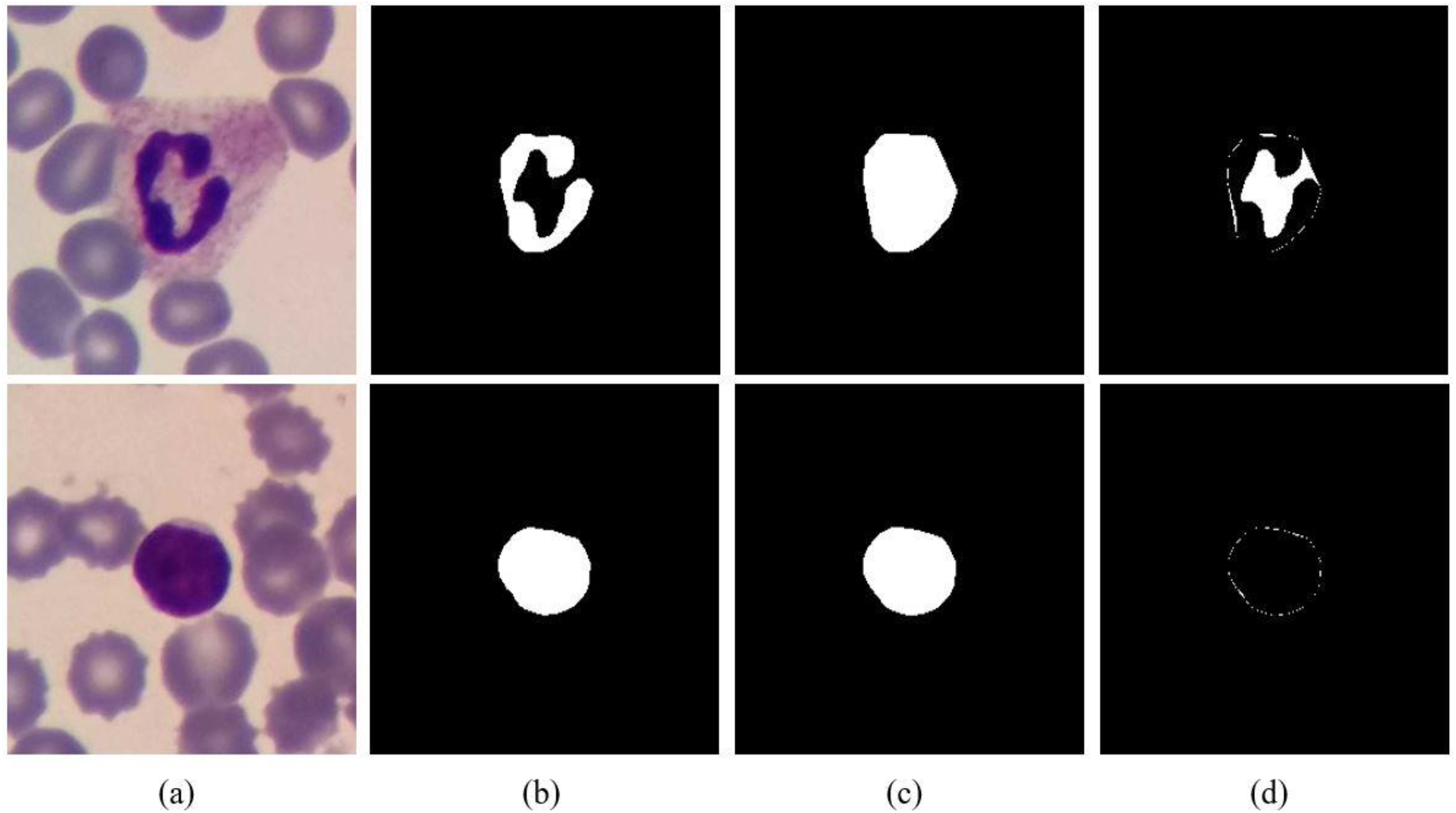
The cytoplasm detection. The first row and the second row are neutrophil and lymphocyte, respectively. (a) RGB image, (b) nucleus, (c) convex hull of the nucleus, (d) the representative of the cytoplasm (ROC).

### Feature Extraction

In this study, two groups of features are taken into account. The first group includes shape features of the nucleus (convexity, circularity, and solidity). The equations associated with the shape features are as follows [1]:

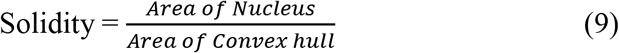

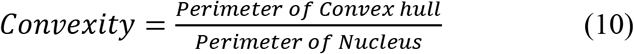

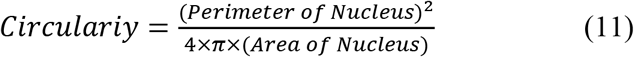

The second group of features is color characteristics. According to the experience of hematologists, in addition to the shape features of the nucleus, the color features of the nucleus and the cytoplasm can also provide us with useful information about the type of WBC [11]. In this research, four novel color features by means of nucleus region, convex hull region, and ROC region are designed as follows:

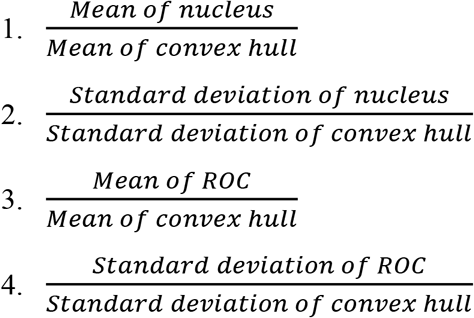

These color features were extracted from the components of RGB, HSV, LAB, and YCrCb color spaces. Therefore, 48 color features and 3 shape features were extracted, which comes up to a total of 51 features. By looking at the classifier’s performance in the results section, it is evident that the introduced color features significantly improve the classification accuracy.

### Classification

After features are extracted from augmented data, they are normalized using the max-min method and are fed into an SVM classifier. We also tested other classifiers such as KNN and deep neural networks. However, we observed that the SVM provides us with the best results. With much trial and error, we found that if the weight of the neutrophils in the training is set to be more than one, and the rest of the classes are one, the best overall accuracy is observed. Three commonly used kernels which are linear, polynomial, and radial basis functions are tested in this regard. Besides, the regularization parameter known as C is an important parameter to train an SVM model. Thus, three important hyperparameters (class-weight, kernel, and C) are tuned to properly train the SVM model. To find the optimal hyperparameters, we applied 5-fold cross-validation on the Train set of the Raabin-WBC employing three different kernels (linear, polynomial with degree three, and radial basis function), neutrophil-weight = 1, 2, 5, 10, 15, 20, and C = 1, 2, 4, 6, 8, 10. Hence, 108 states were assumed. We examined each combination of the hyperparameters with 5-fold cross-validation on the Train set of the Raabin-WBC. Table 2 shows the result of examining different combinations of the hyperparameters. From Table 2, it can be seen that the best accuracy is obtained by polynomial kernel, neutrophil-weight of 10, and this is when the C parameter is equal to 6. We fixed these hyperparameters obtained over the Raabin-WBC dataset meaning that we did not readjust these hyperparameters for the LISC and BCCD datasets.

**Table 2.**
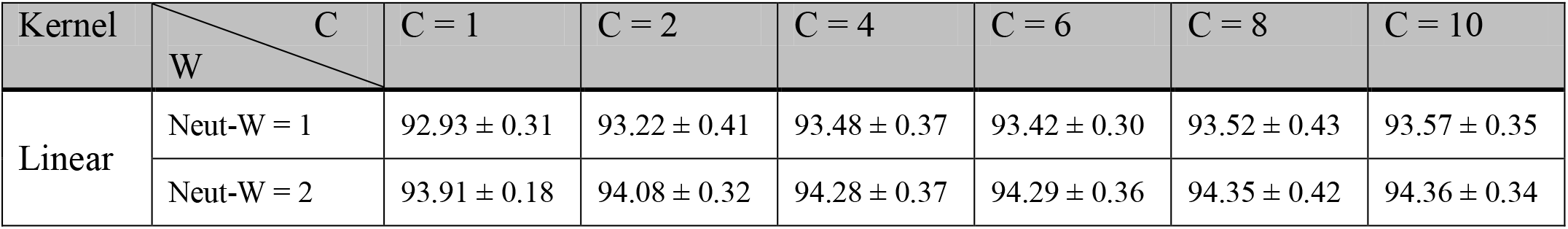
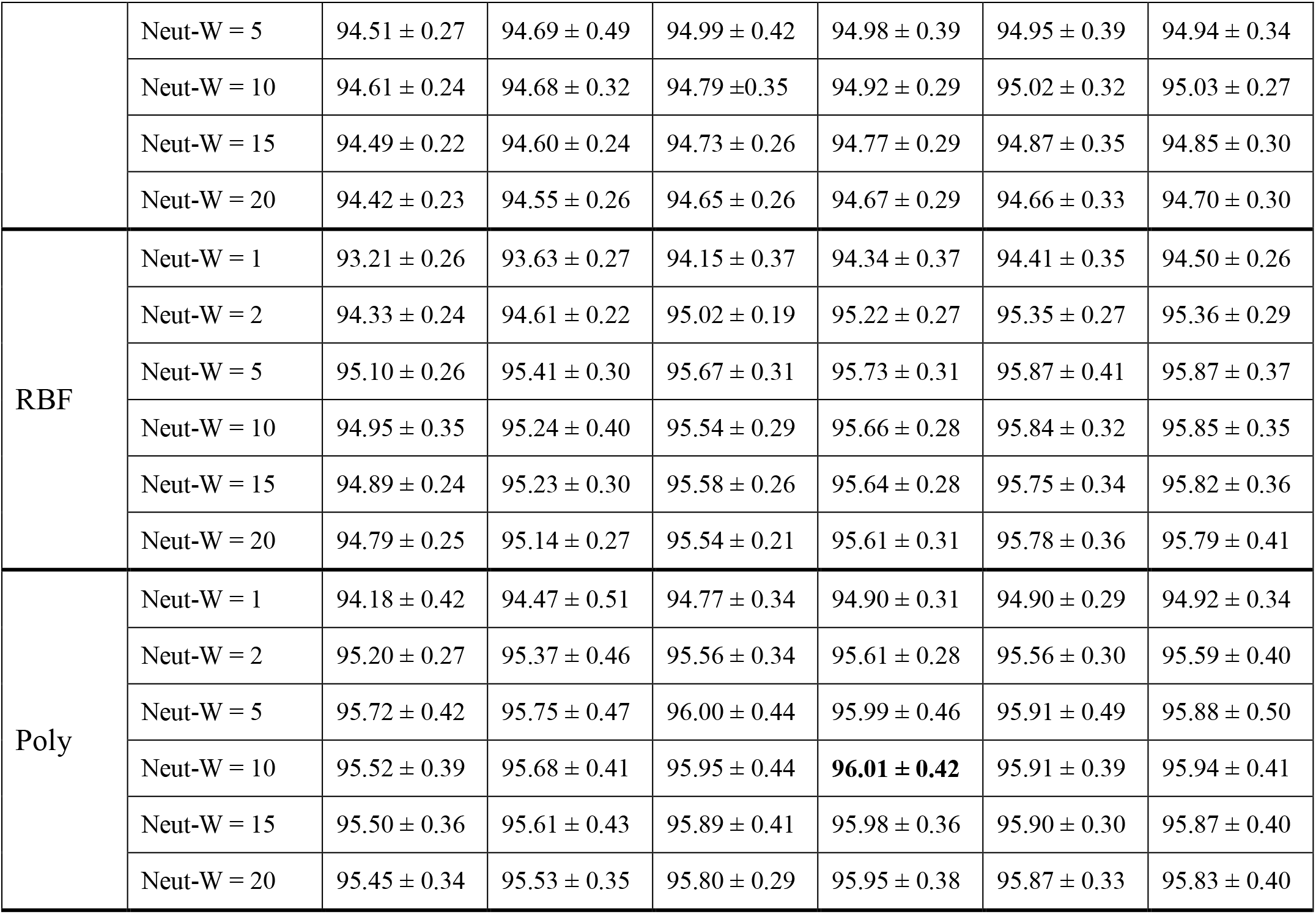
The accuracy for 5-fold cross validation on the Raabin-WBC in order to find the optimal hyperparameters; RBF (radial basis function), Poly (polynomial with degree 3), C (regularization parameter), Neut-W (neutrophil-weight). The results shows that the SVM model with polynomial kernel, C = 6, and neutrophil-weight = 10 provides the best accuracy

## Results

### The Result of Nucleus Segmentation

The performance of the proposed nucleus segmentation algorithm is evaluated using three different metrics namely dice similarity coefficient (DSC), sensitivity, and precision. These metrics are computed using true positive (TP), false positive (FP), true negative (TN) and false negative (FP) of the resulting segmentation (as shown in Figure 5) and are provided by the following equations.

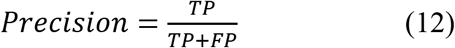

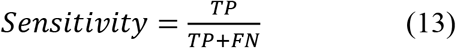

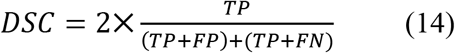

**Figure 5.**
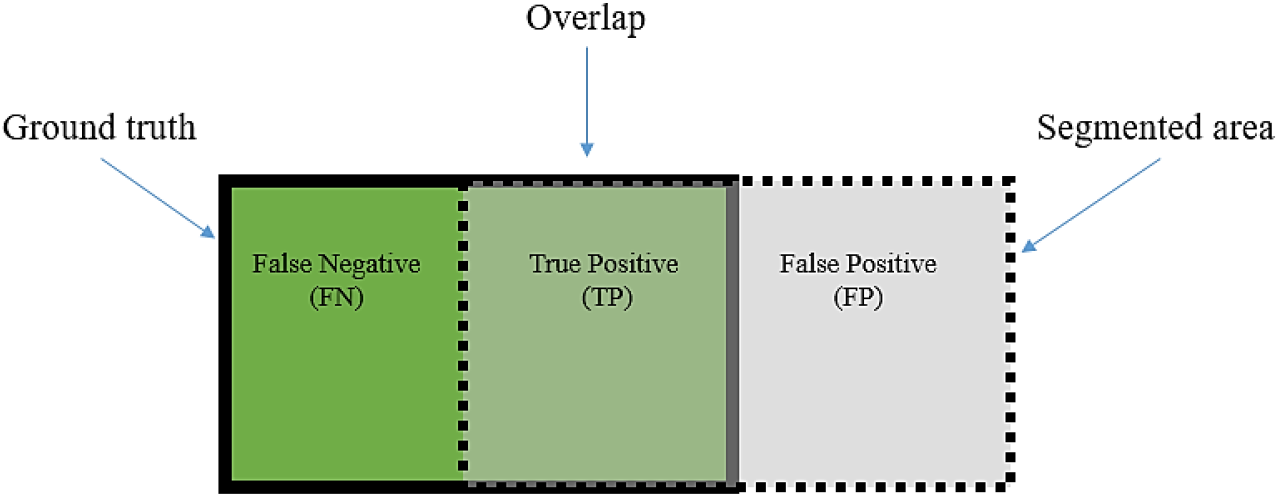
The graphic display of TP, FP, and FN for a segmentation problem.

In order to extract the ground truth, 250 images including 50 images from each type of WBCs were randomly selected from Raabin-WBC dataset. Then, the ground truths for these images were identified by an expert with the help of Easy-GT software [36]. Also, since very dark purple granules cover the basophil’s surface, it is almost impossible to distinguish the nucleus [11]. Therefore, the whole basophil cell was considered as the ground truth. The results of the proposed segmentation algorithm have been presented in Table 3. The proposed segmentation method can detect the nucleus with precision, sensitivity, and dice similarity coefficient of 0.9972, 0.9526, and 0.9675, respectively.

**Table 3.** The result of different nucleus segmentation algorithms evaluated on 250 test images; DSC (dice similarity coefficient), Std (standard deviation), ms (millisecond).

| Methods | Precision |  | Sensitivity |  | DSC |  | Time (ms) | Trainable Parameters |
| --- | --- | --- | --- | --- | --- | --- | --- | --- |
|  | Mean | Std | Mean | Std | Mean | Std |  |  |
| U-Net++ [33] | 0.9598 | 0.0632 | **0.9873** | 0.0225 | **0.9719** | 0.0397 | 1612 | 897412 |
| Attention U-Net [37] | 0.9478 | 0.0903 | 0.9850 | **0.0213** | 0.9633 | 0.0584 | 628 | 854936 |
| Mousavi *et al.* [36] | 0.9362 | 0.1158 | 0.9827 | 0.0310 | 0.9542 | 0.0750 | 47 | 0 |
| Proposed method | **0.9972** | **0.0090** | 0.9526 | 0.0300 | 0.9675 | **0.0180** | **45** | 0 |

The performance of the proposed segmentation algorithm is compared with that of U-Net++ [33], Attention U-Net [37], and Mousavi *et al*.’s method [36]. U-Net++ and Attention U-Net are two well-known deep CNN developed for medical image segmentation. To train these models, 989 images from Raabin-WBC dataset were randomly chosen, and their ground truths were extracted by an expert utilizing Easy-GT software [36]. The training set includes 199 lymphocytes, 199 monocytes, 199 neutrophils, 195 eosinophils, and 197 basophils. Both models were trained for 40 epochs, then evaluated with 250 ground truths mentioned in the previous paragraph. Table 3 presents the results of different segmentation algorithms. It can be seen that the proposed segmentation method has very low standard deviation for DSC and precision which indicates that the proposed method works consistently well for different cells in the data. In addition, U-Net++ and attention U-Net are deep CNNs, and their training process is supervised. Hence, they need way more data to be trained. This is while our proposed method does not need to be learned. Also, these two models have lots of parameters and need more time to segment an image, but the proposed segmentation algorithm is simpler and faster. The suggested method can detect the nucleus of a WBC in a 575 by 575 image size in 45 milliseconds. This is while U-Net++ and attention U-Net need 1612 and 628 milliseconds to segment the nucleus. The proposed method, U-Net++, attention U-Net, and Mousavi *et al*.’s method [36] were implemented in Google Colab, CPU mode and were compared their execution time.

### Result of Classification

In order to evaluate the classification accuracy, four metrics are used: Precision, Sensitivity, F1-score (F1), and Accuracy (Acc). If we face a two-class classification problem such the first class is called Positive and the second class is called Negative, the confusion matrix can be assumed as Table 4, and the mentioned criteria are obtained through relations (15), (16), (17), and (18).

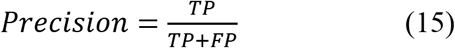

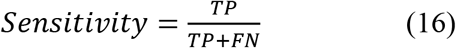

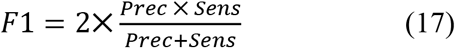

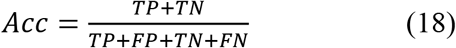

**Table 4.** Confusion matrix for a two-class problem.

|  | Positive | Negative |
| --- | --- | --- |
| Positive | **True Positive (TP)**<br>Number of samples that are Positive and classifier predicts Positive | **False Negative (FN)**<br>Number of samples that are Positive while classifier predicts Negative |
| Negative | **False Positive (FP)**<br>Number of samples that are Negative while classifier predicts Positive | **True Negative (TN)**<br>Number of samples that are Negative and classifier predicts Negative |

In order to evaluate the effectiveness of color features, Raabin-WBC, LISC, and BCCD datasets are classified in two modes: classification using the shape features, and classification using the shape features together with the color ones. The comparison of the classification accuracy of these two modes is provided in Table 5. It can be seen in Table 5 that adding proposed color features significantly changes the classification results. Addition of color features leads to a remarkable increase in precision, sensitivity, and F1-score for all five types of WBCs. The proposed method classifies WBCs in Raabin-WBC, LISC, and BCCD datasets with accuracies of 94.65 %, 92.21 %, and 94.20 %, respectively. The resulting confusion matrices of our proposed method for the three datasets are shown in Figure 6.

**Table 5.** The comparison of classification results using two modes of features. One mode uses only shape features and the other mode uses both shape and color features. The abbreviations: P (precision), S (sensitivity), F1 (F1-score), Acc (accuracy).

| Dataset | Features | Lymph. |  |  | Mon. |  |  | Neut. |  |  | Eos. |  |  | Bas. |  |  | Acc. (%) |
| --- | --- | --- | --- | --- | --- | --- | --- | --- | --- | --- | --- | --- | --- | --- | --- | --- | --- |
|  |  | P (%) | S (%) | F1 (%) | P (%) | S (%) | F1 (%) | P (%) | S (%) | F1 (%) | P (%) | S (%) | F1 (%) | P (%) | S (%) | F1 (%) |  |
| Raabin-WBC | Shape | 95.63 | 93.04 | 94.31 | 45.13 | 37.61 | 41.03 | 83.82 | **96.99** | 89.93 | 0 | 0 | 0 | 65.00 | 43.82 | 52.35 | 84.56 |
|  | Shape & Color | **97.23** | **95.07** | **96.14** | **84.87** | **86.32** | **85.59** | **98** | 95.60 | **96.78** | **72.24** | **91.30** | **80.66** | **96.59** | **95.51** | **96.05** | **94.65** |
| LISC | Shape | 82.35 | 77.78 | 80 | 81.82 | 60 | 69.23 | 62.96 | **100** | 77.27 | 28.57 | 18.18 | 22.22 | 53.33 | 50.00 | 51.61 | 64.94 |
|  | Shape & Color | **84.21** | **88.89** | **86.49** | **84.62** | **73.33** | **78.57** | **94.44** | **100** | **97.14** | **100** | **100** | **100** | **100** | **100** | **100** | **92.21** |
| BCCD | Shape | 0 | **0** | 0 | 0 | 0 | 0 | 71.07 | 100 | 83.05 | 0 | 0 | 0 | --- | --- | --- | 71.01 |
|  | Shape & Color | **100** | **100** | **100** | **100** | **100** | **100** | **94.12** | **97.96** | **96** | **88.89** | **72.73** | **80** | --- | --- | --- | **94.20** |

**Figure 6.**
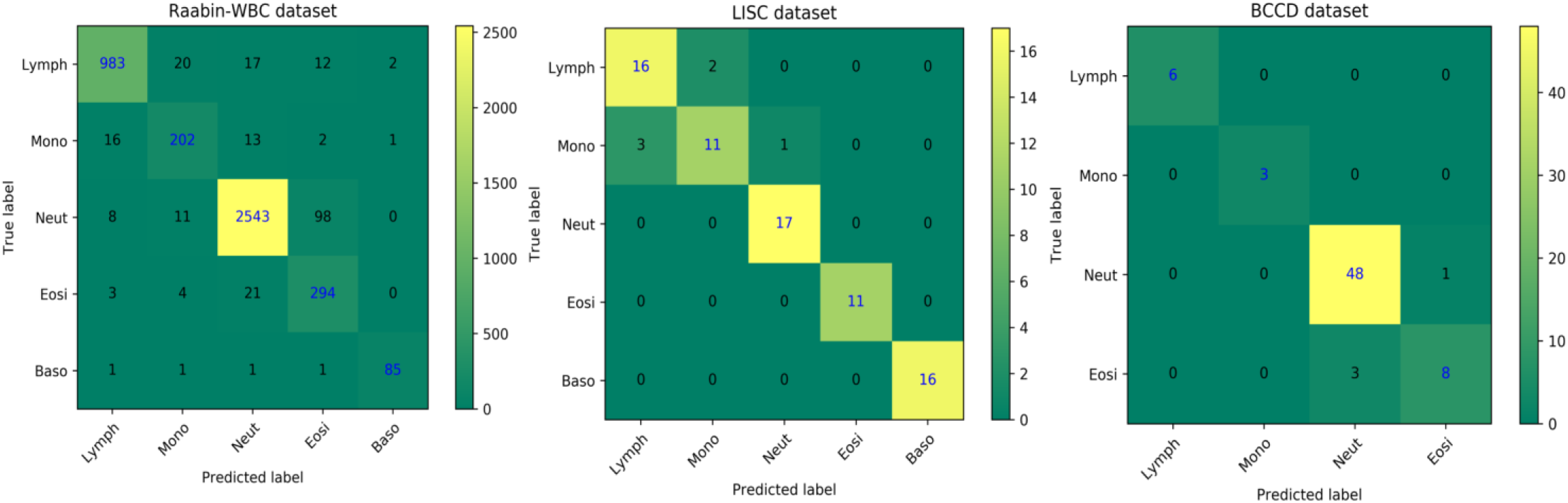
The confusion matrices of our proposed classification method for Raabin-WBC, LISC, and BCCD datasets.

### Comparison with the state-of-the-art methods

Since the LISC and BCCD datasets have been publicly available for several years, the performance of the proposed method on these two datasets is compared to that of the state-of-the-art works in terms of precision, sensitivity, and F1-score. Also, because the categorization of WBCs in peripheral blood is an imbalanced classification problem [15], the comparison has been made based on each class. Table 6 shows the detailed comparisons.

**Table 6.** The comparison of our method with other works; P (precision), S (sensitivity), F1 (F1-score).

| Dataset | Method | Lymph. |  |  | Mon. |  |  | Neut. |  |  | Eos. |  |  | Bas. |  |  |
| --- | --- | --- | --- | --- | --- | --- | --- | --- | --- | --- | --- | --- | --- | --- | --- | --- |
|  |  | P (%) | S (%) | F1 (%) | P (%) | S (%) | F1 (%) | P (%) | S (%) | F1 (%) | P (%) | S (%) | F1 (%) | P (%) | S (%) | F1 (%) |
| LISC | Rezatofghi and Soltanian-Zadeh [7] | 100 | 93.10 | 96.43 | 92 | 95.83 | 93.88 | 93.33 | 100 | 96.55 | 100 | 94.74 | 97.30 | 90.74 | 98 | 94.23 |
|  | Jung *et al.* [22] | 100 | 88.89 | 94.12 | 100 | 66.67 | 80 | 94.44 | 100 | 97.14 | 100 | 100 | 100 | 73.91 | 100 | 85 |
|  | Baydilli and Atila [23] | 100 | 100 | 100 | 80 | 80 | 80 | 91.66 | 100 | 96.65 | 100 | 83 | 90.71 | 88.89 | 88.89 | 88.89 |
|  | **Our method** | 84.21 | 88.89 | 86.49 | 84.62 | 73.33 | 78.57 | 94.44 | 100 | 97.14 | 100 | 100 | 100 | 100 | 100 | 100 |
| BCCD | Banik *et al.* [24] | 99 | 100 | 99.50 | 100 | 81 | 89.50 | 74 | 97 | 83.95 | 96 | 84 | 89.60 | --- | --- | --- |
|  | Liang *et al.* [25] | 100 | 100 | 100 | 96 | 80 | 87.27 | 78 | 92 | 84.42 | 93 | 91 | 91.99 | --- | --- | --- |
|  | Banik *et al.* [26] | 100 | 100 | 100 | 99 | 99 | 99 | 93 | 92 | 92.50 | 93 | 93 | 93 | --- | --- | --- |
|  | Togacar *et al.* [19] | 97.99 | 99.80 | 98.89 | 95.83 | 100 | 97.87 | 93.49 | 92.09 | 92.78 | 94.92 | 90.86 | 92.85 | --- | --- | --- |
|  | **Our method** | 100 | 100 | 100 | 100 | 100 | 100 | 94.12 | 97.96 | 96 | 88.89 | 72.73 | 80 | --- | --- | --- |

By taking a meticulous look at criterion F1-score, which actually covers both criteria precision and sensitivity, it can be said that our proposed method has achieved the best performance in most classes. In the LISC dataset, the proposed method has classified neutrophils, eosinophils, and basophils with F1-scores of 97.14%, 100%, and 100%, respectively. Also, in the BCCD dataset, our method was able to classify lymphocytes, monocytes, and neutrophils with F1-scores of 100%, 100%, and 96%, respectively. In reference to traditional approaches, the method employed in this article is simple and creative and can be easily implemented. In this method, suitable shape and color features are extracted by means of the nucleus and the cytoplasm, yet there is no need for the cytoplasm to be segmented. The methods used in [19], [22], [23], [24], [25], and [26] are based on deep learning approaches. Therefore, their models are more complex and have more trainable parameters versus our classifier model which is SVM. For example, the models utilized in [22], [23], and [25] have 16.5, 23.5, and 59.5 million parameters, successively. Besides, it should be noted that the hyperparameters of our SVM model were set only using the Raabin-WBC dataset and were not readjusted again on the LISC and BCCD datasets. This is while the other methods have fixed the hyperparameters of their classifiers on each dataset, separately.

## Discussion

As mentioned before, the proposed method contains three phases. Segmenting the nucleus and detecting a part of the cytoplasm located in the nucleus’s convex hull are performed at first phase. After extracting shape and color features, WBCs are finally categorized employing extracted features. Our proposed nucleus segmentation algorithm consists of several steps depicted in Figure 2. These steps have been designed to remove the red blood cells and the cytoplasm. From Table 3, it is clear that the segmentation algorithm can detect the nucleus with a very high precision of 0.9972 and DSC of 0.9675. The proposed segmentation algorithm is very fast in comparison with U-Net++ and Attention U-Net models (Table 2).

In the cytoplasm detection phase, in contrast to the common practice of segmenting the whole cytoplasm, only parts of the cytoplasm that are inside the convex hull of the nucleus was selected as a representative of cytoplasm (ROC). This way has not the difficulties of segmenting cytoplasm, but the classification accuracy is boosted with the help of features extracted by means of ROC.

In the Feature extracting phase, we used three common shape features namely solidity, convexity, and circularity. Besides, we designed four novel color features and extracted them from channels of RGB, HSV, LAB, and YCrCb color spaces. According to Table 5, it is obvious that the designed color features have remarkably increased the classification accuracy.

In the final phase, the classification is done with an SVM model. To choose the best hyperparameters for the SVM model, 5-fold cross validation was applied only on the Raabin-WBC dataset. The SVM model was separately trained for a different combination of hyperparameters to obtain the best one (Table 2). The method we put forward is automatic and simple that does not need to resize the images and segment the cytoplasm. According to Table 6, in LISC dataset, the proposed method came first in distinguishing neutrophils, eosinophils, and basophils. In addition, in the BCCD data set, our method was ranked first in detecting lymphocytes, monocytes, and neutrophils.

## Conclusion

In this research, we designed a novel nucleus segmentation algorithm and four new color features to classify WBCs. The proposed method successfully managed to classify three data sets differing in terms of microscope, camera, staining technique, variation, and lighting conditions, and ensured the following accuracy of 94.65 % (Raabin-WBC), 92.21 % (LISC), and 94.20 % (BCCD). We showed that the novel color features designed for this research can greatly help identifying WBCs. Therefore, we can conclude that not only is the suggested method robust and reliable, but also it can be utilized for laboratory applications and purposes.

## Data and code availability

Three used dataset are available through the below links:

- Raabin-WBC: www.raabindata.com
- LISC: http://users.cecs.anu.edu.au/~hrezatofighi/Data/Leukocyte%20Data.htm
- BCCD: https://www.kaggle.com/paultimothymooney/blood-cells

Also, the codes of the proposed method are available from first author on reasonable request.

## Author contributions

S. T. implemented the proposed segmentation and feature extraction algorithms and wrote the paper. Z. M. K. suggested the feature extraction algorithm and helped the first author in writing the paper as well as implemented U-Net and Attention U-Net models. A. G. and R. H. revised the manuscript and made some comments and suggestions. A. G. designed the research as supervisor. All authors reviewed the final version of the manuscript before submitting.

## Additional information

**Correspondence** and requests for materials should be addressed to E.T.

